# A Single Kinetic Parameter Determines Intermediate Filament Elongation

**DOI:** 10.64898/2026.11.20.768144

**Authors:** Lovis Schween, Harald Herrmann, Norbert Mücke, Ben Fabry, Stéphanie Portet

## Abstract

Intermediate filaments elongate by end-to-end annealing of filament subunits. We develop a Smoluchowski coagulation model with a single kinetic parameter, *k*_0_, that reproduces data from time-resolved atomic force microscopy, electron microscopy, and static light scattering. The model predicts lognormal filament length distributions, sublinear filament growth, and saturation of scattering intensity ratios. For a given protein, the inferred assembly rate constants agree across measurement techniques.

---

Intermediate filaments (IFs) are a major component of the cytoskeletal network in metazoan cells. Their proteins constitute a multigene family whose members are expressed in a tissue-specific manner. For example, vimentin is predominantly found in mesenchymal cells, whereas desmin is characteristic of muscle cells. Both proteins assemble into extended, approximately 10 nm-wide filaments that contribute to the mechanical resilience of cells [1].

To investigate IF assembly kinetics, purified proteins are initially maintained in low-salt buffers, where they form soluble tetrameric complexes. A rapid increase in ionic strength triggers the lateral association of typically eight to twelve tetramers into approximately 60 nm-long structures, termed unit-length filaments (ULFs). This initial assembly step is completed within about one second. ULFs define the elementary units of filament length, while subsequent growth proceeds through binary end-to-end annealing of filaments of different lengths [2, 3]. Owing to longitudinal overlap, each incorporated ULF contributes approximately 45 nm to the contour length of the growing filament.

The resulting filament length distributions can be determined by electron microscopy (EM) and atomic force microscopy (AFM) at successive assembly times, typically spanning seconds to minutes [4–6]. Static light scattering, by contrast, resolves assembly kinetics on millisecond timescales but does not directly yield filament length distributions. Combining these complementary approaches therefore offers a means to connect ensemble scattering signals with the underlying kinetics of filament elongation.

In this letter, we combine the slow but single-filament-resolved microscopy methods with fast but ensemble-averaged dual wavelength light scattering (DWLS) method. We exploit two predictions from a theoretical framework for IF assembly that requires only a single kinetic parameter, *k*_0_, to quantify the filament elongation.

First, we derive that the filament length distribution is described by a log-normal distribution, and second, that the growth of the geometric mean filament length follows a power-law in time, with exponents slightly below unity. We then experimentally demonstrate the universality of both theoretical predictions using microscopy and light scattering data under a variety of assembly conditions and for different IF types. Finally, we present a method to compute the kinetic parameter *k*_0_ regardless of the measurement technique.

## Mechanistic model

We describe IF elongation by the Smoluchowski coagulation equation. Filaments of length *i* ULFs (*i ≥* 1) have time-dependent concentrations *F*_*i*_(*t*) and elongate exclusively by end-to-end annealing. Their dynamics follows

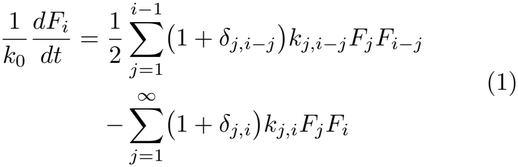

The first term describes the formation of a filament of length *i* from two shorter ones, the second its loss by annealing with any other filament. We assume that filament fragmentation is negligible during the first 30 minutes of assembly so that no breakage terms are included [7]. We numerically integrate this equation for initial condition *F*_1_(0) *>* 0 (initial concentration *c*_0_ of ULFs), and *F*_*i>*1_(0) = 0 (initially only ULFs are present).

The diffusion-limited rate constants *k*_*i,j*_ = *k*_*j,i*_ are calculated using semiflexible polymer theory [8, 9]. It accounts for center-of-mass diffusion and internal bending fluctuations that bring filament ends into contact [10]. Hence, *k*_*i,j*_ depend on the contour length of both engaging filaments and on the persistence length *l*_*p*_, with *l*_*p*_ *≈* 1 *µ*m for both vimentin and desmin. The kinetic parameter *k*_0_ is dimensionless and captures the biochemical specificity of reactive ends; it is the only free parameter of the mechanistic model.

Equation (1) exhibits a time-rescaling symmetry: solutions for arbitrary (*c*_0_, *k*_0_) follow from reference solutions computed at 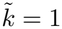 and 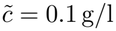 via

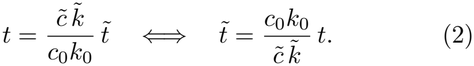

Any observable *O*(*t*; *k*_0_, *c*_0_) thus satisfies the time-rescaling relation 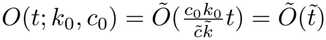.

Equation (1) predicts the concentrations of the complete filament population, *F*_*i*_(*t*), but experimental techniques do not report *F*_*i*_(*t*) directly. Microscopy counts individual filaments and samples the number-weighted length distribution, while light scattering weights every filament by its squared mass and its wavelength-dependent form factor. Since both observables originate from the same *F*_*i*_(*t*) and therefore from the same *k*_0_, they are interchangeable. Below, we express the observables of both techniques as functions of *k*_0_.

## Observables

Microscopy yields the length fraction distribution of the filaments

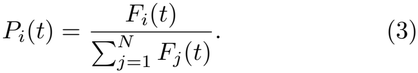

and geometric mean filament length *m*_*g*_(*t*)

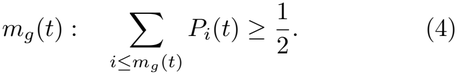

For static light scattering, the scattered intensity *I* at wavelength *w* is proportional to the mass-weighted average of the squared form factor 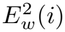, with 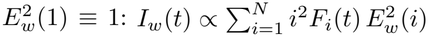.

The dual-wavelength intensity ratio, our central observable, is

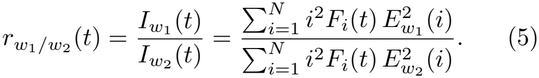

Following Rayleigh-Gans theory, 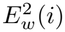 decreases more rapidly at shorter wavelengths as *i* increases [11]. Hence, the ratio rises monotonically as the mean filament length increases over time. As filaments grow beyond about 6-7 ULFs the ratio becomes insensitive to further elongation and saturates at a plateau set by the ratio of the Rayleigh–Gans form factors at wavelengths *w*_1_ and *w*_2_ [11, 12]. To account for experimental variations mainly due to baseline drift between measurements, we introduce one additional parameter *f* when we fit the model to the DWLS observable, 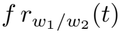, Fig. 2(A).

**FIG 1.**
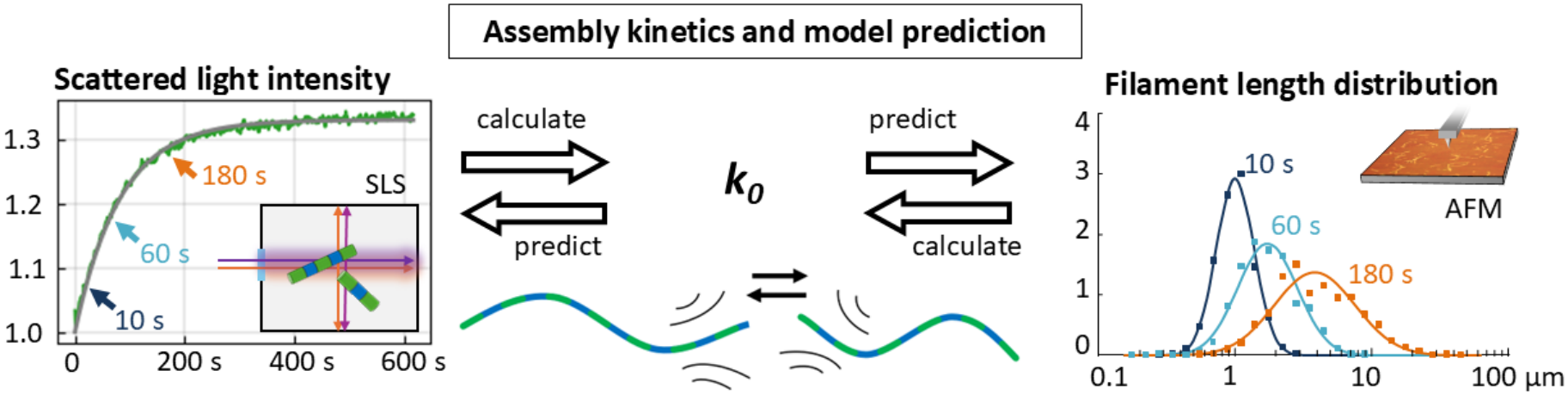
Overview of the unified approach to investigate the elongation of intermediate filaments as semiflexible biopolymers. The common kinetic parameter *k*_0_ connects the *in vitro* elongation kinetics recorded by static light scattering (left) and microscopy (right). Furthermore, this framework enables the *k*_0_-based prediction of measurement results. Time points (10 s, 60 s, 180 s) of the filament length distributions are indicated in the DWLS intensity ratio curve.

**FIG 2.**
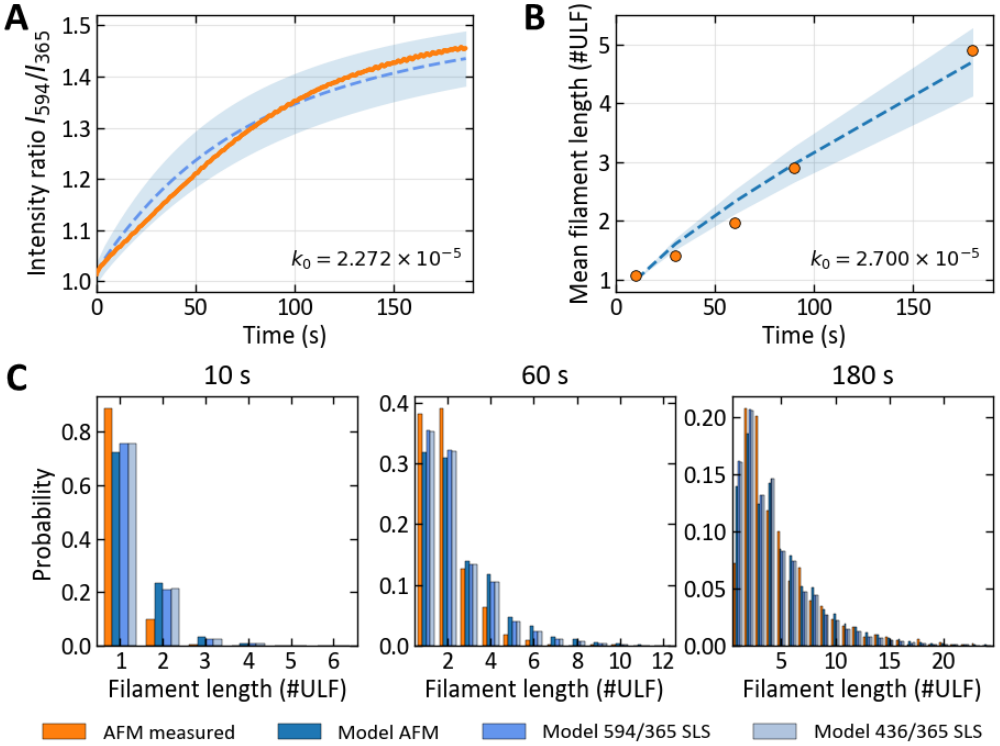
Mechanistic-model fits and cross-modal predictions. (A) Fit to the DWLS intensity ratio *I*_594_*/I*_365_ over 180 s. (B) Fit to the AFM-derived mean filament length in units of ULFs. Dashed blue curves denote best fits, shaded regions the 95% confidence bands, and orange symbols the measurements. (C) AFM filament-length distributions at 10, 60, and 180 s (orange bars) compared with predictions based on *k*_0_ inferred independently from AFM, *I*_594_*/I*_365_, or *I*_436_*/I*_365_ (blue).

## Model fit

Fitting Eq. (1) to AFM-derived mean lengths yields *k*_0_ = 2.7 *×* 10^*−*5^, Fig. 2(B), while an independent fit to the DWLS ratio gives *k*_0_ = 2.3 *×* 10^*−*5^, Fig. 2(A), with overlapping confidence intervals. With either value, the model correctly predicts the measured length distributions at 10, 60 and 180 s without further adjustment, Fig. 2(C). The mechanistic model thus unifies both techniques, but each fit requires repeated numerical integration, which is time-consuming. We therefore approximate the model predictions using *k*_0_ dependent theory-informed algebraic expressions.

## Algebraic form of model predictions

From model reduction and numerical integration, three laws emerge. First, the numerically obtained length distribution *L*(*t*) closely follows a log-normal distribution during the first 30 min of assembly

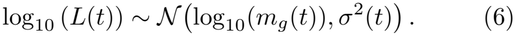

Second, the distribution width *σ*(*t*) is directly linked to the geometric mean length *m*_*g*_(*t*) of the filaments, approximately following the relationship

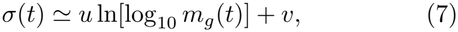

with *u ≈* 0.055 *±* (3 *×* 10^*−*4^) and *v ≈* 0.33 *±* (2 *×* 10^*−*4^). Eq. (7) applies for both persistence lengths considered (*l*_*p*_ = 0.3 and 1 *µ*m), however, it cannot be extrapolated toward the monodisperse initial state: at *m*_*g*_ = 1, the relation is undefined.

Third, *m*_*g*_(*t*) grows in time according to a power law

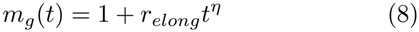

with

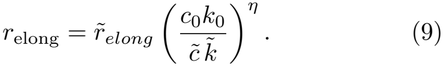

Fitting Eq. (8) to Eq. (4) under reference conditions (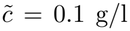 and 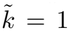) yields *η ≈* 0.89 *±* (6 *×* 10^*−*5^), independent of *l*_*p*_, and 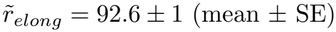.

Solving Eq. (9) for *k*_0_ yields

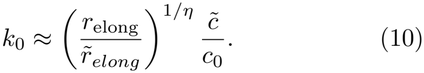

Hence, Eq. (10) directly links the geometric mean growth coefficient *r*_elong_ to *k*_0_ for a known protein concentration *c*_0_, Fig. 3 (A).

**FIG 3.**
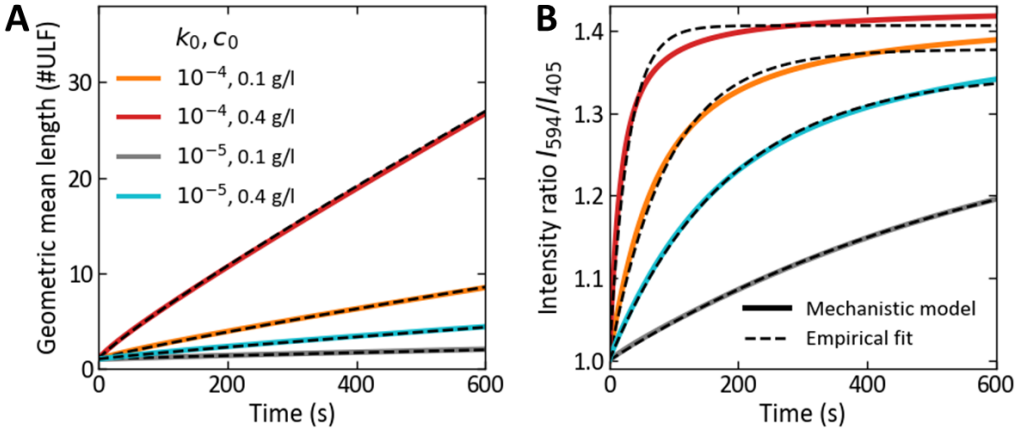
Empirical models. (A) Fitted lognormal geometric mean *m*_*g*_ (*t*) = 10^*µ*(*t*)^ (plain curves) compared with the sub-linear growth Eq. (8) (dashed curves) for reaction parameters *k*_0_ = 10^*−*4^ and 10^*−*5^ and protein concentrations *c*_0_ = 0.1 and 0.4 g/l. (B) Predicted intensity ratios *I*_594*nm*_*/I*_405*nm*_ (plain curves) for the same (*k*_0_, *c*_0_) pairs shown in (A), following Eq. (5) with filament *l*_*p*_ = 1 *µ*m. Fitted saturating exponential model Eq. (11) (dashed curves).

For DWLS experiments, an analogous compact description of intensity ratios *r*(*t*) is provided by a saturating exponential model (Fig. 3B):

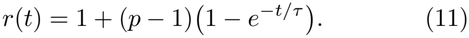

Here, *τ* is a time constant that describes how fast the ratio approaches the plateau value *p*. Therefore, *τ* provides a quantitative measure of elongation kinetics.

The theoretical plateau 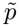 is set by the Rayleigh-Gans form factor and has a value of 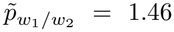 for *w*_1_*/w*_2_ = 594*/*405 used in our experiments. Under reference conditions 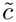 and 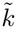, we find numerically 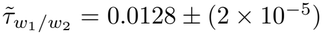 for *w*_1_*/w*_2_ = 594*/*405. For any (*c*_0_, *k*_0_) pairs, omitting wavelength dependency for readability, *τ* follows

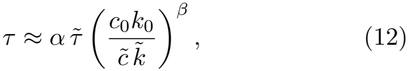

with coefficients *α* = 2.13 *±* 0.4 and *β* = *−*0.87 *±* 0.02 for *w*_1_*/w*_2_ = 594*/*405.

Solving Eq. (12) for *k*_0_ yields

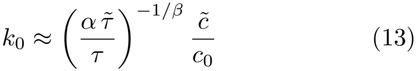

Hence, Eq. (13) directly links the time constant for the intensity ratio *τ* to *k*_0_ for a known protein concentration *c*_0_, Fig. 3 (B). The values of *k*_0_ determined with either of the empirical approximations (Eqs. (13) or (10)) closely agree with those obtained by full optimization of the mechanistic model while reducing the required computation time from hours to milliseconds.

## Cross-modal prediction

The experimentally measured parameters *r*_elong_ and *τ* can be converted into one another by equating Eqs. (9) and (12):

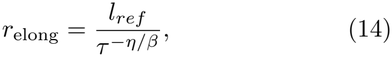

with reference length 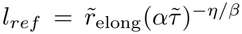 in units of ULFs. Because the exponent satisfies *−η/β ≈* 1.02, Eq. (14) reduces to

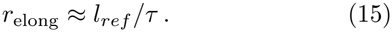

Thus, Eqs. (14) and (15) connect the DWLS relaxation time constant *τ* to the growth coefficient of the geometric mean filament length *r*_*elong*_.

## Validation

To assess the consistency between the two measurement techniques, we compare *k*_0_-values obtained with DWLS and AFM for vimentin and desmin, preferably under matched assembly conditions (shaded area in (Fig. 4 A), for protein concentrations *c*_0_ = 0.1–0.4 g*/*l, salt concentrations of 50, 100, and 160 mM, and in NaPi, Tris-HCl, or MOPS buffers [12–14]. The paired estimates of *k*_0_ obtained with both techniques lie close to the line of identity (Fig. 4 B), demonstrating that consistent *k*_0_ values are obtained, independently of the measurement modality. Moreover, under otherwise identical assembly conditions, *k*_0_ remains approximately independent of protein concentration, consistent with concentration affecting the assembly timescale through the product *c*_0_*k*_0_ (Eq. (2))

**FIG 4.**
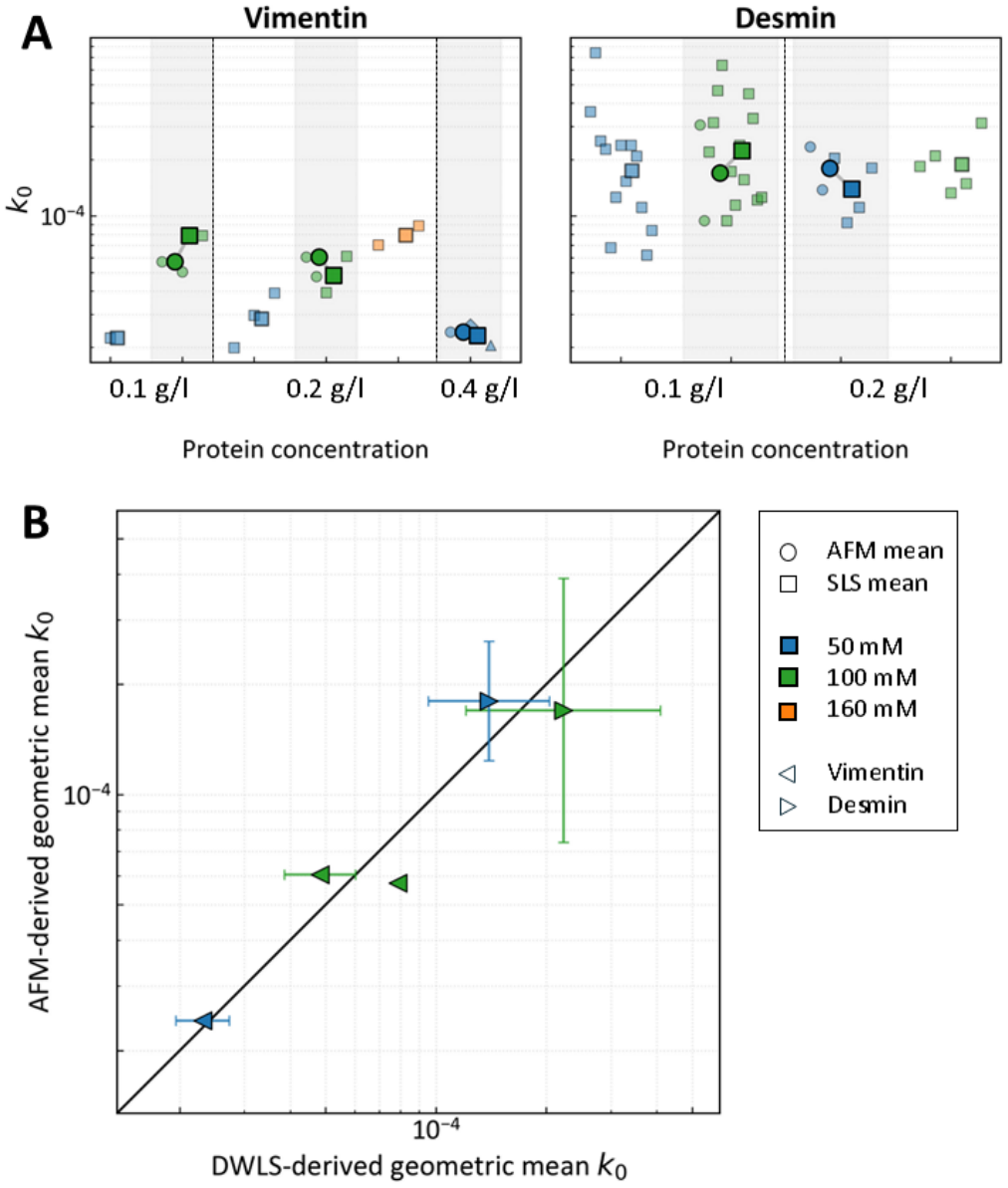
(A) Kinetic parameter *k*_0_ derived from DWLS time constant *τ* of the exponential fit to the ratio *w*_1_*/w*_2_ = 594*/*405 (squares) and the AFM-derived median growth coefficient *r*_elong_ (circles) for vimentin (left, *<*) and desmin (right, *>*) filaments, after assembly of 0.1, 0.2 or 0.4 g/l protein at 50 mM (blue), 100 mM (green) or 160 mM (orange). Bigger, opaque symbols show mean of *k*_0_ of the respective assembly condition and method. Grey shaded area marks measurements from both methods at same assembly conditions. (B) Correlation of AFM-derived and DWLS-derived *k*_0_ values at the same assembly conditions (shaded in A).

Comparing *k*_0_ across assembly conditions reveals several robust trends, Fig. 4 (A). First, *k*_0_ increases approximately linearly with ionic strength in the case of vimentin in Fig. 4 (A), which is consistent with electrostatic screening of the reactive filament ends. Desmin assembly, however, is considerably less salt dependent. Second, within the investigated conditions, no systematic dependence on the buffer is found for either protein.

Third, under equivalent buffer conditions, *k*_0_ is approximately threefold higher for desmin than for vimentin, in agreement with the faster desmin assembly reported previously [14].

The algebraic form of model predictions also carry physical information. According to Eq. (9), the growth coefficient scales as *r*_elong_ *∝* (*c*_0_*k*_0_)^*η*^: protein concentration and end reactivity enter only through their product, whereas the exponent *η* reflects how the encounter rate depends on filament size. If all filaments annealed at the same rate regardless of their length, the mean length would grow linearly in time, *η* = 1. More generally, dynamic scaling analysis of the Smoluchowski equation shows that an association kernel homogeneous of degree *λ, k*_*ci,cj*_ = *c*^*λ*^*k*_*i,j*_, yields an asymptotic mean size that grows as *t*^1*/*(1*−λ*)^ [15]. A kernel that decreases with size (*λ <* 0) therefore produces sublinear growth. Identifying *η* = 1*/*(1 *λ*) gives *λ ≈ −*0.12, hence the effective association rate decreases only weakly with filament length. This compensation becomes stronger with filament size: the effective homogeneity changes from *λ ≈ −*0.10 for encounters between short filaments to *λ ≈ −*0.08 for mixed pairs and *λ ≈ −*0.05 for encounters between long filaments, as obtained from a numerical analysis of the semiflexible association kernel *k*_*i,j*_ in Eq. (1). The association kinetics therefore becomes practically size-independent as the filaments grow.

This vanishing size dependence reflects a competition characteristic of semiflexible polymers. As filaments elongate, hydrodynamic drag reduces their translational and rotational mobility, whereas thermal bending fluctuations of their reactive ends enhance the search for an annealing partner [8, 9]. The near-zero value of *λ* indicates that these local fluctuations largely compensate for the loss of global filament mobility. This interpretation is consistent with scaling theory and Brownian-dynamics simulations predicting an approximately length-independent annealing rate when transverse end fluctuations dominate filament encounters [16].

The framework reported in this letter is restricted to the elongation-dominated phase during the first 30 min of assembly. At longer times, fragmentation and subunit exchange along the filament contour become important and would require extending Eq. (1) to include breakage and recombination. Moreover, microscopy becomes unreliable for very long filaments, whereas the DWLS intensity ratio loses sensitivity as it approaches its form-factor-determined plateau. But apart from this experimental limitation, the underlying strategy - exploiting the time-rescaling symmetry of the coagulation equation and approximating its observables in closed form - applies wherever coagulation kinetics are probed with complementary real-space and ensemble techniques.

Physically, the nearly size-independent association kernel shows that local fluctuations of filament ends can off-set the global hydrodynamic slowing of growing semiflexible polymers, giving rise to an unexpected simplification of IF assembly that is governed by a single rate constant *k*_0_. Thus, *k*_0_ provides a common quantitative measure of IF elongation kinetics, enabling systematic comparisons across proteins, ionic conditions, and molecular perturbations. Relating these kinetic differences to molecular structure may reveal the common physical principles underlying filament elongation across the diverse IF protein family.

S.P. is supported by the Natural Sciences and Engineering Research Council of Canada (RGPIN-2025-06063). L.S. gratefully acknowledges funding of the Er-langen Graduate School in Advanced Optical Technologies (SAOT) by the Bavarian State Ministry for Science and Art.

## Notes

### Competing Interest Statement

The authors have declared no competing interest.

